# The central nucleus of the amygdala and gustatory cortex assess affective valence during CTA learning and expression in male and female rats

**DOI:** 10.1101/2021.05.04.442519

**Authors:** Alyssa Bernanke, Elizabeth Burnette, Justine Murphy, Nathaniel Hernandez, Sara Zimmerman, Q. David Walker, Rylee Wander, Samantha Sette, Zackery Reavis, Reynold Francis, Christopher Armstrong, Mary-Louise Risher, Cynthia Kuhn

## Abstract

This study evaluated behavior (Boost® intake, LiCl-induced behaviors, ultrasonic vocalizations (USVs), task performance) and c-Fos activation during conditioned taste aversion (CTA), the reinforced task (Boost® task) and control task (cage only) to understand how male and female rats balance the relative danger or safety of a stimulus in learning and performing a task. Females drank more Boost® than males but showed similar aversive behaviors after LiCl treatment. Males produced 55 kHz USVs (indicating positive valence) when anticipating Boost® and inhibited these calls after pairing with LiCl. Females produced 55 kHz USVs based on their estrous cycle but were more likely to make 22 kHz USVs than males (indicating negative valence) after pairing with LiCl. c-Fos responses were similar in males and females after Boost® or LiCl. Females engaged the gustatory cortex and ventral tegmental area more than males during the Boost® task and males engaged the amygdala more than females in both the reinforcing and devalued tasks. Network analysis of correlated c-Fos responses across brain regions identified two unique networks characterizing the Boost® and LiCl (CTA) tasks, in both of which the VTA played a central role. RNAscope identified a population of D1-receptor expressing cells in the CeA that responded to Boost® and D2 receptor-expressing cells that responded to LiCl. The present study suggests that males and females differentially process the affective valence of a stimulus to produce the same goal-directed behavior.

## Introduction

The ability to properly evaluate the relative safety of a stimulus is critical to survival across species. Dysfunction of this mechanism contributes to a range of mental health disorders, including anxiety disorders such as generalized anxiety, obsessive compulsive disorders, and post-traumatic stress disorders ^1–5^. Anxiety disorders are one of the most common neuropsychiatric disorders, with a lifetime prevalence of 28.8% ^6^. Perturbations in the ability to assess the relative danger or safety of a situation is a key component that drives anxiety disorders.

Interrogation of the neural circuits that mediate normal and maladaptive safety evaluation have relied on fear conditioning paradigms, in which a neutral cue is paired with an aversive stimulus to produce a conditioned response ^7–9^. These fear-learning circuits offer important insight into the formation and retrieval of fear memory ^10–13^. However, these paradigms are limited in their ability to address how an animal balances the safety and/or danger of a stimulus, a process that more closely recapitulates real-world decision-making.

Here we use conditioned taste aversion (CTA) to model the complexity of this decision-making process. CTA is a classical conditioning paradigm which pairs a palatable substance with an aversive visceral experience, such as the emetic agent lithium chloride (LiCl), producing an aversion to the substance at subsequent exposure. Unlike many other classical conditioning paradigms, in which an aversive stimulus is paired with a neutral cue, the use of a palatable substance in CTA requires the animal to process the affective valence of the reward versus the aversion. This provides an opportunity to understand neural processes by which this decision-making occurs.

We used several modifications of typical published approaches to better capture the ability of this task to assess affective valence. We used a nutritive substance (chocolate-flavored Boost®) in non-food deprived animals, which more closely represents CTA outside the confines of a laboratory ^14^. While current studies of CTA focus primarily on performance of the task, this study aimed to capture both the emotional valence and the neural mechanisms that respond to each step of the CTA process. We accomplished this goal by evaluating both behavior and c-Fos responses to the appetitive stimulus (Boost®), the aversive stimulus (LiCl), and the performance of the CTA task. In the control condition, animals are simply placed in the control environment, compared with performance of the task when Boost® paired with NaCl, and so are anticipating a reinforcing stimulus only, and the traditional task in which Boost® is initially paired with LiCl. By including the acute conditions, as well as the control and appetitive task, we were better able to capture previously underappreciated neural activation in brain regions relevant to the CTA paradigm.

We included females in the present study. Females are disproportionately affected by anxiety-related disorders, including post-traumatic stress disorder, obsessive-compulsive disorder, generalized anxiety, and phobias ^5, 15–19^. Despite these known sex differences, female rodents are critically understudied in animal models of neuropsychiatric disorders including CTA, of which only 2% of studies utilize female rodents ^20^. We aimed to determine if any sex differences in the CTA response reflected sex differences in responses to the aversive stimulus, palatable stimulus, or recall and performance of the task.

In this study, we characterized behavioral and neural responses to CTA in males and females. Behavioral observations of the aversive stimulus included pica, ptosis, and lying on belly (LOB) as indications of nausea in LiCl-treated rats. We also quantitated ultrasonic vocalizations (USVs) as a measure of emotional valence on task day. Rats engage in two distinct call patterns: short 55-kHz (which range from 30-70 kHz and are associated with positive emotional valence), and longer 22-kHz calls (associated with negative emotional valence). They emit 55 kHz calls during behaviors such as play, mating, social approach, and in anticipation of a reward ^21–24^. We therefore hypothesized that animals would engage in these calls on task day when anticipating Boost® that was not devalued with LiCl injection. Conversely, 22-kHz calls are exhibited in contexts of social withdrawal (e.g. after mating ^25^) and to warn the colony of potential predators ^26–28^. Female rats are more likely than males to exhibit 22-kHz calls in the presence of cat urine ^29^. We hypothesized that rats would engage in these calls on task day when anticipating Boost® that was previously paired with LiCl.

We used the immediate early gene c-Fos as a marker of neuronal activity at each stage of CTA: in response to the acute rewarding, aversive, and neutral stimuli (NaCl injection), and during control task, Boost® task and during CTA expression. We analyzed 11 brain regions known to be associated with the response to reward and/or LiCl. We selected more rostral regions that are associated with the decision-making process in CTA, rather than the more caudal regions which serve to transmit sensory signals that are interpreted by higher-order areas of the brain. We also conducted network analysis to interrogate correlations among the c-Fos responses to the areas sampled. Finally, our c-Fos data prompted us to examine more closely specific cell types in the amygdala using RNAScope, a novel mRNA in situ hybridization technique.

The following studies showed that males and females differentially assign valence to the expectation of a rewarding and aversive stimulus, identified cell types that contribute to the neural processing of these stimuli, and characterized novel behaviors that correlate with predicted CTA expression and experimental condition.

## Results

### Behavioral responses during conditioned taste aversion

The CTA experimental design is described in Figure 1a. To first characterize CTA expression, a dose-response curve was established in both sexes. Animals received either NaCl (0.15M) or LiCl (19 mg/kg, 38 mg/kg, or 80 mg/kg) (Figure 1b). LiCl-induced behavior was measured as described in Table 3 in Materials and Methods. Statistics show a main effect of sex [F (1,111) =6.05, p=0.015] and a main effect of treatment [F (3,111) =29.31, p< 0.001] but no interaction of treatment x sex. Fisher’s post hoc analysis showed that males inhibited Boost® intake more than females after pairing with the low dose of 19 mg/kg LiCl.

**Figure 1.**
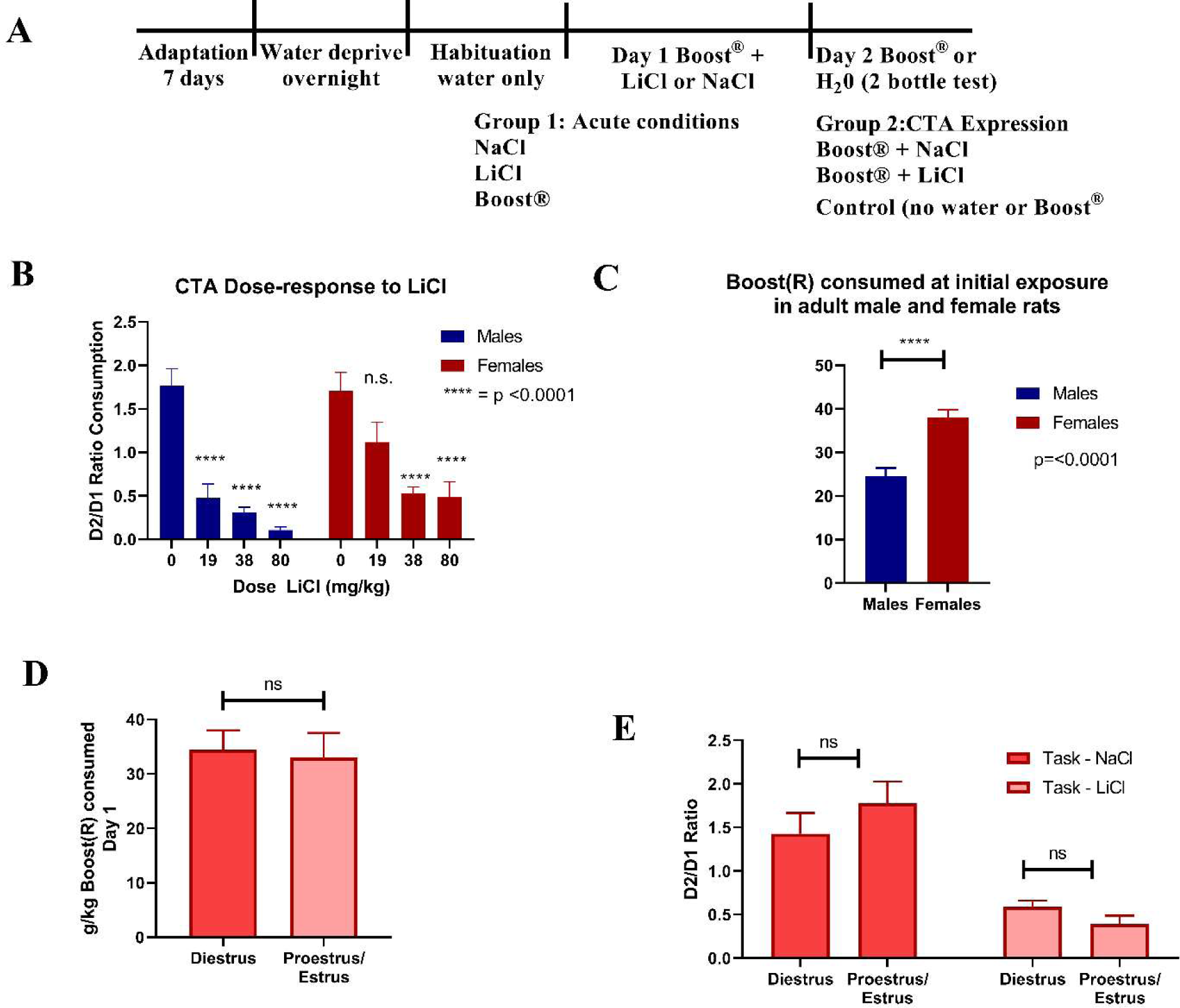
(A) Protocol for CTA paradigm. (B) Dose response curve to LiCl (n=7-23). (C) Boost® consumption on first exposure by sex (n=48-100). (D) Estrous cycle and day 1 Boost® consumption (n=8- 9). (E) Estrous cycle and CTA expression on task day (n=10-14). All data expressed as mean±SEM.

Female rats drank more Boost® than males at initial exposure when adjusting for body weight, with a main effect of sex [F (1,146) =21.91, p< 0.001, (Fig. 1c)]. Both male and female rats generally increased the volume of Boost® they consumed at subsequent exposure. This ratio of consumption was similar in males and females despite the propensity for females to drink more on first exposure.

To determine whether estrous cycle contributed to CTA behavior, estrous cycle was determined by vaginal lavage on task day. Cycle stage did not have a significant effect on CTA expression (Fig. 1d).

### LiCl-induced behaviors

Pica was measured after the same range of doses of LiCl: 0 mg/kg (NaCl), 19, 38, and 80 mg/kg. Both male and female rats engaged in pica equally across doses of LiCl. 2-way ANOVA showed an effect of treatment [F (3,158) =20.79, p< 0.001] and no effect of sex or interaction of treatment x sex (Fig. 2a). Pica events at 80 mg/kg were higher than 38 and 19 mg/kg. Lying on belly (LOB) was the only nausea behavior measured in this study that correlated with CTA expression, Pearson correlation = -0.3701 and p < 0.0434 (Fig. 2b, c). Male and female rats also exhibited ptosis comparably after LiCl injection (Fig. 2d). Neither of these behaviors correlated with CTA behavior as measured by the ratio of Boost® consumption on day 2 as compared to day 1.

**Figure 2.**
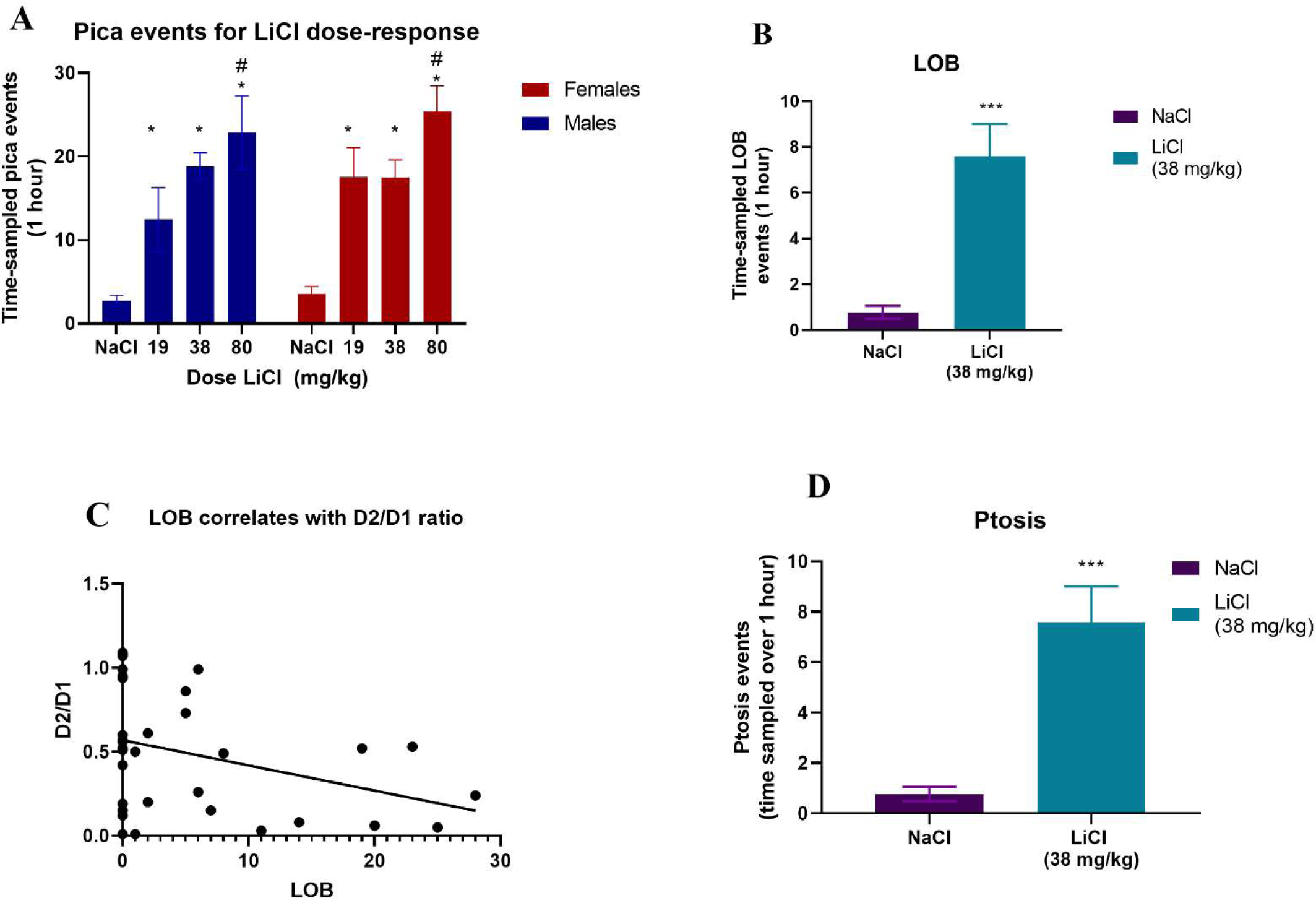
(A) Pica events after increasing doses of LiCl (0, 19, 38, and 80 mg/kg) in male and female rats (n=6-45) (B) LOB was measured for 1 hour after 38 mg/kg LiCl or NaCl (n=18-33). Data shown as mean ± SEM. (C) LOB correlation with D2/D1 ratio (D) Ptosis events measured over 1 hour after 38 mg/kg LiCl or NaCl (n=18-33). All data shown as mean±SEM. *=different from NaCl (0 mg/kg). # = different from 19 mg/kg and 38 mg/kg. *** = p<0.0005.

### Ultrasonic vocalizations during conditioned taste aversion task

Rats were taken through the CTA protocol as described. We recorded USVs during the first 10 minutes of the habituation process on Day 2, before offering the 2-bottle test, as vocalizations typically occur in anticipation of a reward.

We found an effect of treatment [F (1,59) =4.52, p< 0.038, Fig. 3a-b], with Boost® task showing more 55 kHz calls than the LiCl task. Dividing by sex due to pre-planned contrast showed that males expressed more 55 kHz calls in the Boost® task [F (1,24) =8.53, p=0.007], while females were not significant.

**Figure 3.**
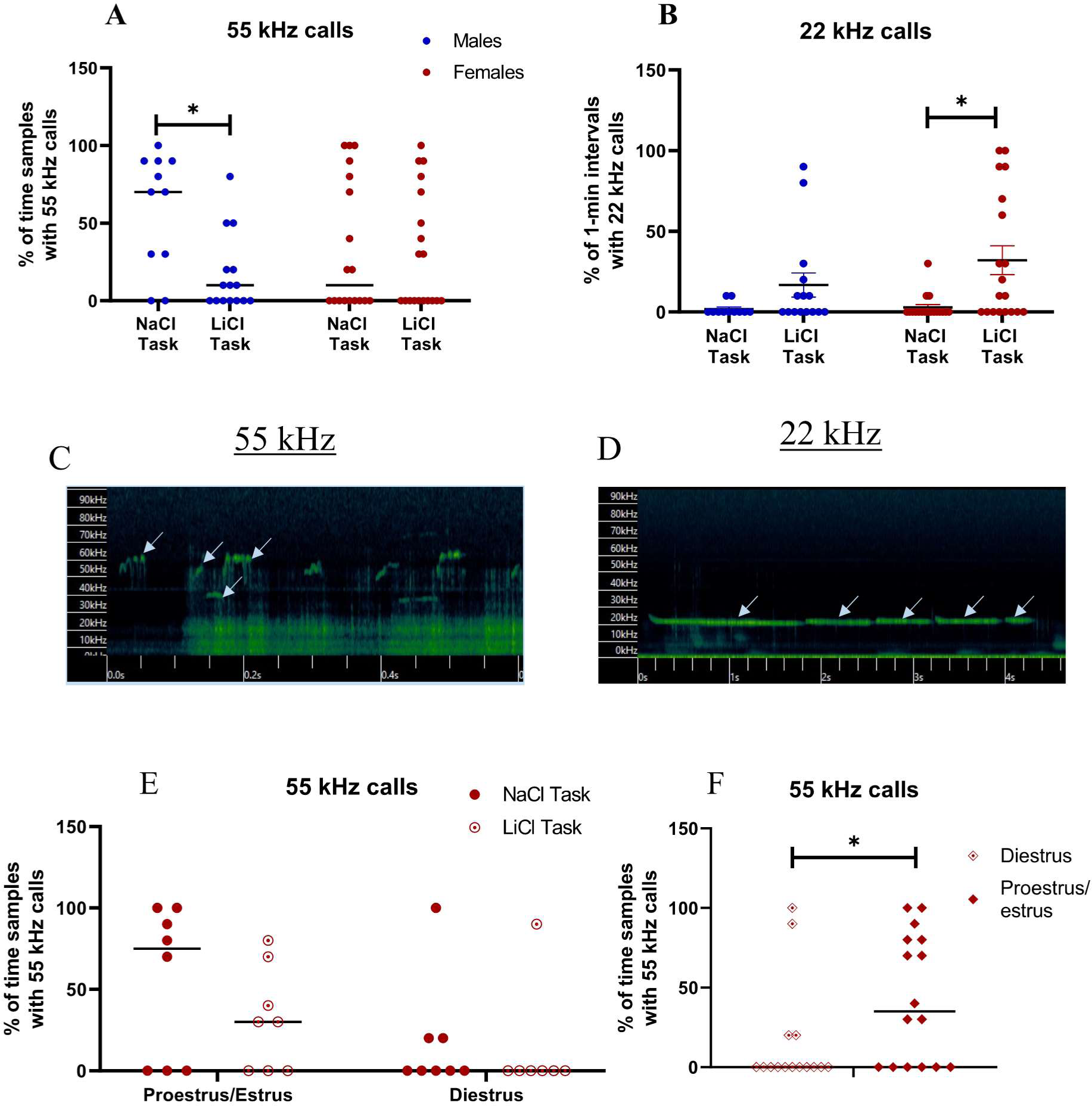
USVs were recorded for first 10 minutes during CTA task. (A) 55 kHz calls in male and female rats during CTA expression (n=11-19). (B) Representative 55 kHz call. (C) 22 kHz warning calls during CTA expression in male and female rats (n=12-19). (D) Representative 22 kHz call (E) 55 kHz vocalizations in females by cycle only (n=150-16) (E) 55 kHz calls in females by task and cycle (n=6-8). Data expressed as mean±SEM. *indicates different by Fisher’s post hoc except where otherwise

There was a significant correlation of 22 kHz calls in the LiCl-paired environment by 2- way ANOVA, with a main effect treatment [F (1,59) =9.48, p< 0.0032, Fig. 3c-d]. Dividing by sex due to pre-planned contrast showed that females significantly increased 22 kHz calls in the LiCl-paired environment [F (1,34) =9.17, p< 0.005], while males were not significant. This effect was not influenced by estrous cycle state (data not shown).

Female rat 55 kHz vocalizations were substantially influenced by estrous cycle, with high-estradiol states associated with an increased likelihood of engaging in 55 kHz calls, and low-estradiol states decreasing the frequency of 55 kHz calls, regardless of experimental condition [F (1,27) =6.35, p < 0.018, Fig. 3e]. By task, females showed an effect of estrous cycle with no effect of treatment [F (1,24) =7.38, p< 0.012, Fig. 3f]. One-way ANOVA for each low- and high-estradiol states individually showed females in high-estradiol states increased 55 kHz vocalizations in the Boost® task compared to LiCl task [F (1,12) =5.20, p< 0.042, Fig. 3f]. There was no effect of treatment in the low-estradiol group.

### The c-Fos response to acute and conditioned stimuli

We evaluated c-Fos expression to better characterize differences in the neural circuits involved in the development and expression of the control, Boost® and LiCl task (CTA) in males and females. c-Fos analysis was performed across 11 brain regions associated with the response to rewarding and aversive stimuli (Table 1).

**Table 1.** c-Fos positive neurons±SEM

|  | Acute NaCl |  | Acute Boost® |  | Acute LiCl |  | Control Task |  | Boost® Task |  | LiCl Task |  |
| --- | --- | --- | --- | --- | --- | --- | --- | --- | --- | --- | --- | --- |
|  | F | M | F | M | F | M | F | M | F | M | F | M |
| vmPFC | 244±37 | 229±72 | 244±20 | 239±33 | 266±41 | 272±51 | 246±37 | 241±32 | 268±52 | 212±31 | 249±44 | 221±43 |
| aIC | 183±32 | 187±40 | 198±20 | 215±22 | 212±50 | 206±27 | 232±32 | 193±14 | 223±31 | 176±26 | 227±21 | 174±24 |
| gIC5/6 | 100±38 | 87±44 | 197±29 | 178±77 | 206±44 | 206±70 | 81±45 | 86±26 | 162±53 | 145±23 | 144±55 | 99±72 |
| gICL4 | 134±66 | 85±28 | 262±23 | 253±82 | 255±23 | 267±89 | 148±82 | 98±64 | 225±69 | 184±27 | 121±45 | 79±43 |
| NAcC | 91±30 | 98±35 | 133±7 | 98±20 | 100±62 | 109±29 | 110±1 | 76±18 | 123±26 | 99±34 | 101±21 | 64±29 |
| NAcS | 117±39 | 123±36 | 125±19 | 107±33 | 171±78 | 141±68 | 120±26 | 90±17 | 153±38 | 113±41 | 124±52 | 118±52 |
| SON | 7±5 | 14±8 | 11±9 | 10±12 | 112±34 | 112±3 | 10±10 | 2±1 | 21±6 | 27±27 | 5±2 | 45±70 |
| BLA | 34±8 | 32±8 | 56±16 | 53±8 | 67±25 | 58±15 | 52±19 | 24±5 | 61±10 | 42±3 | 52±6 | 41±15 |
| CeA | 44±22 | 42±26 | 133±40 | 108±33 | 191±73 | 199±58 | 30±7 | 42±60 | 142±19 | 115±60 | 62±48 | 48±28 |
| PVN | 113±51 | 103±26 | 122±51 | 143±29 | 230±102 | 226±96 | 95±36 | 107±28 | 91±20 | 74±45 | 141±39 | 134±70 |
| VTA | 63±17 | 50±13 | 84±30 | 92±16 | 93±29 | 52±15 | 63±22 | 69±15 | 118±22 | 66±30 | 83±13 | 51±9 |

### c-Fos immunoreactivity during CTA acquisition and expression

We performed 3-way repeated measures ANOVA (sex x treatment, area as a repeated measure) on all experimental conditions, which revealed a main effect of sex [F (1,62) =5.37, p=<0.001, females greater than males], a main effect of treatment [F (5,62) =9.57, p=<0.01], a main effect of area [F (10,462) =229.69, p <0.001], and an interaction of treatment x area [F (50,462) =8.46, p< 0.001].

We then performed 3-way repeated measures ANOVA for the acute and conditioned stimuli separately. For the acute condition, 3-way repeated measures ANOVA (treatment x sex, area as a repeated measure) revealed a main effect of treatment [F (2,31) =17.36, p < 0.001] and area [F (10,231) =101.64, p < 0.001], and an interaction of treatment x area [F (20,231) = 9.73, p < 0.001], with no effect of sex for the acute stimuli. Acute Boost® and acute LiCl groups both differ from NaCl control, and different from each other (LiCl greater than Boost®). We conducted lower level 2-way ANOVA (sex x Rx) for areas that are implicated functionally in CTA and/or showed statistically relevant findings.

The central nucleus of the amygdala (CeA) responds to aversive visceral stimuli such as LiCl as well as rewarding stimuli ^30^. We found c-Fos in the CeA increased both to Boost® and LiCl [F (2,30) = 31.39, p< 0.001 effect of treatment] similarly in male and female rats (Fig. 4a, atlas location shown in Fig. 4c, representative images shown in Fig. 4d). The basolateral amygdala (BLA), known for its role in assessing the valence of a stimulus ^31–34^, showed an effect of treatment [F (2,17) = 9.06, p< 0.0021], with no effect of sex (Fig. 4b, atlas location shown in Fig. 4c, representative images shown in Fig. 4e). Post-hoc analysis showed both Boost® and LiCl groups were increased compared to NaCl control.

**Figure 4.**
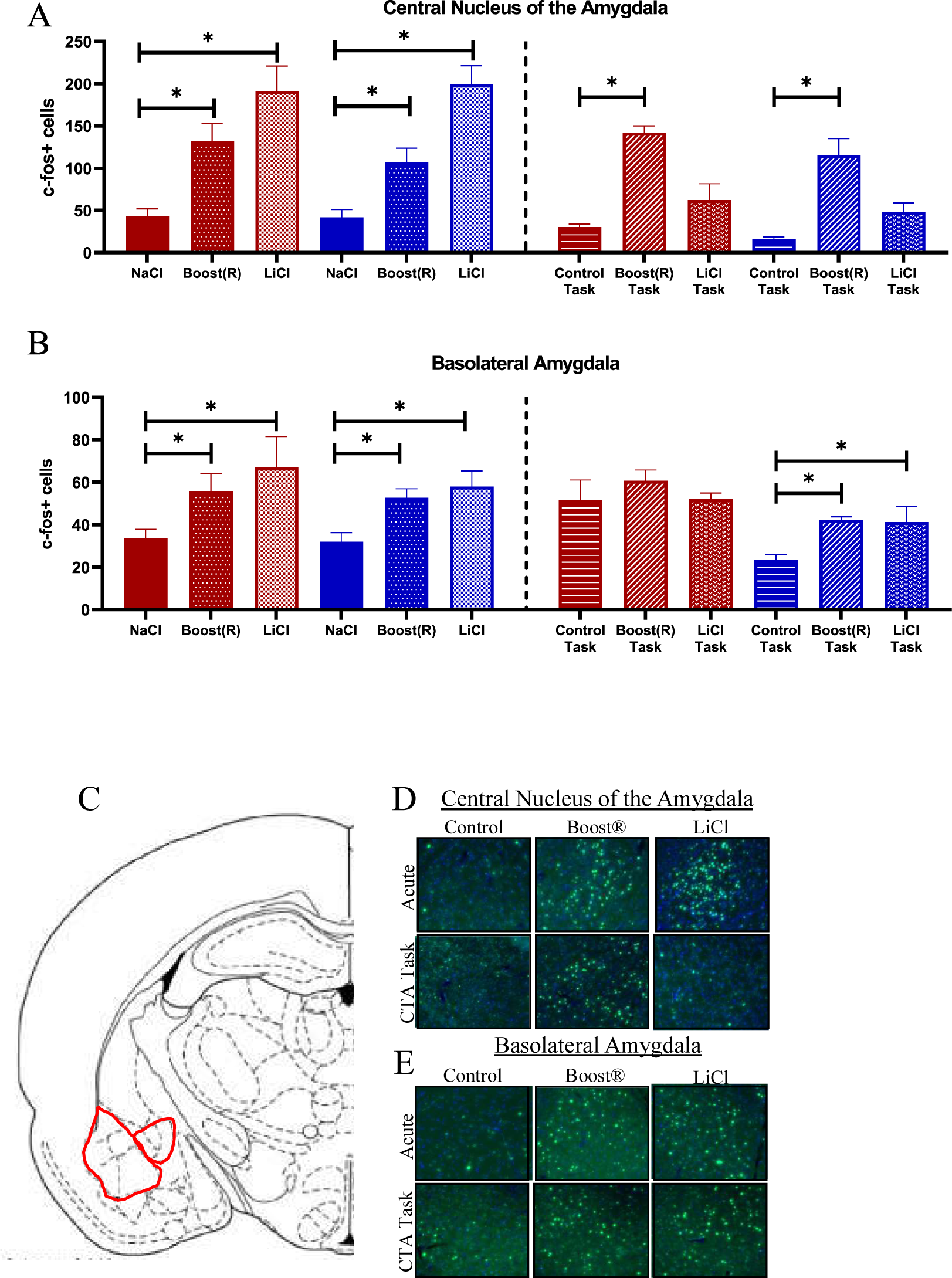
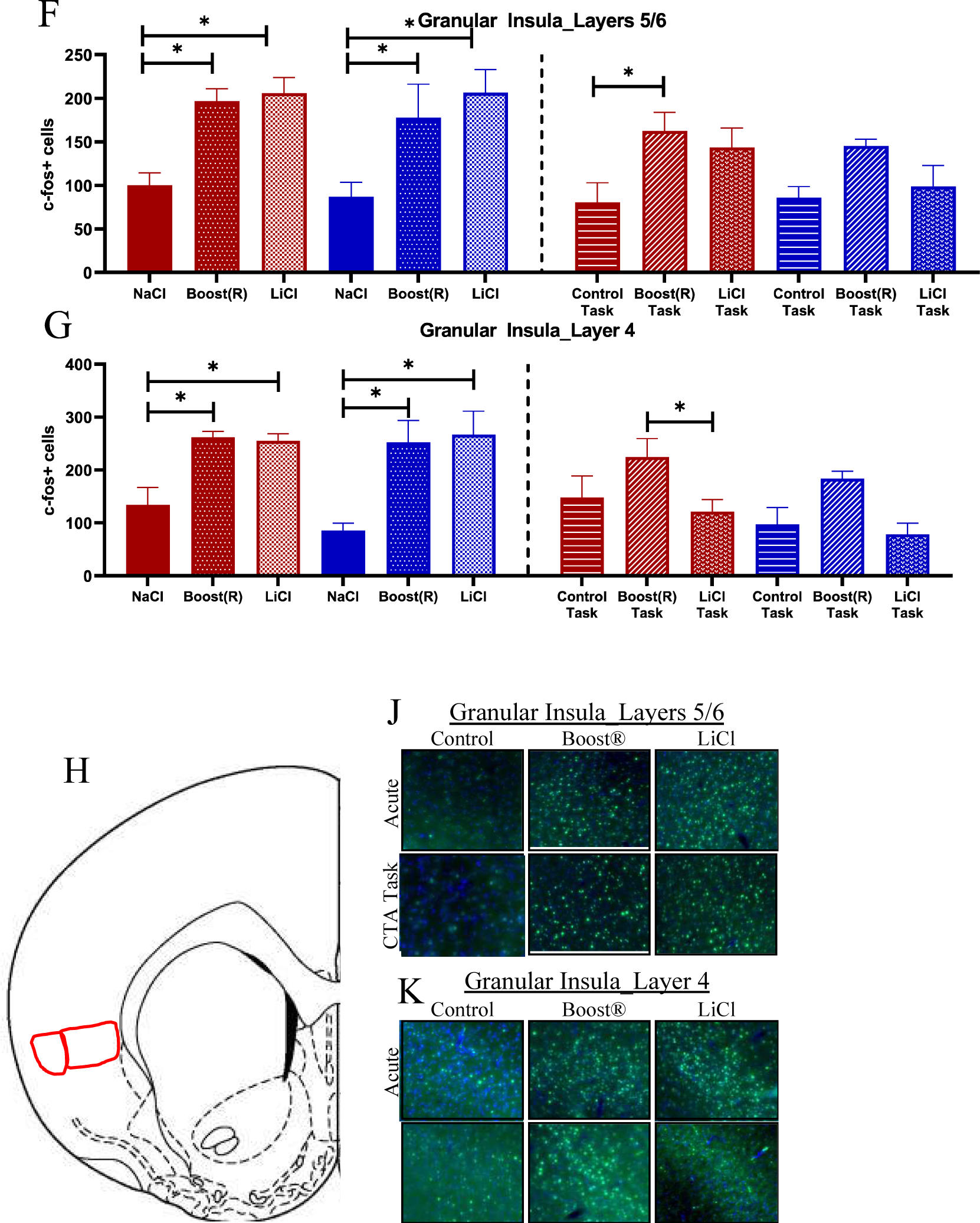
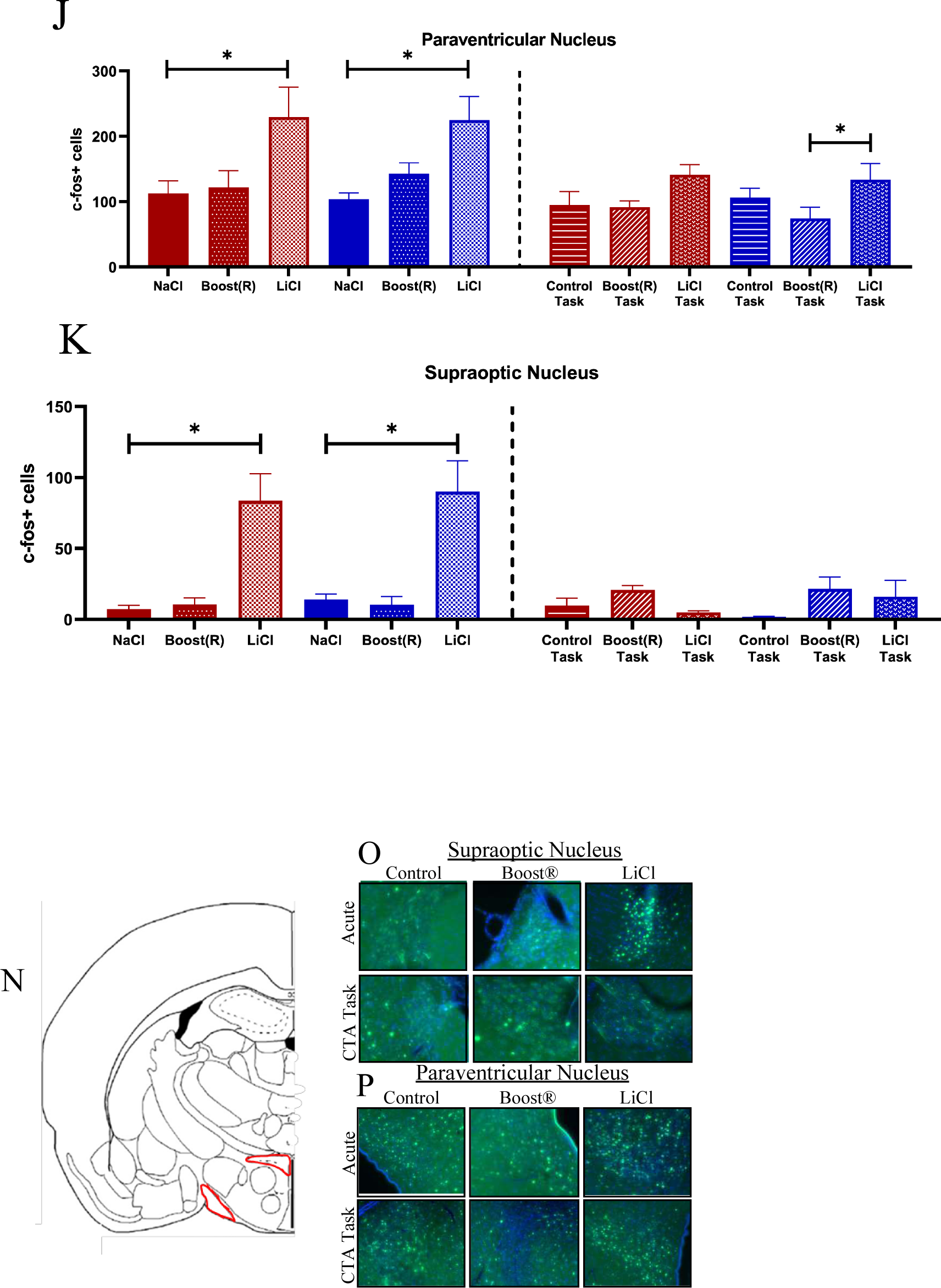
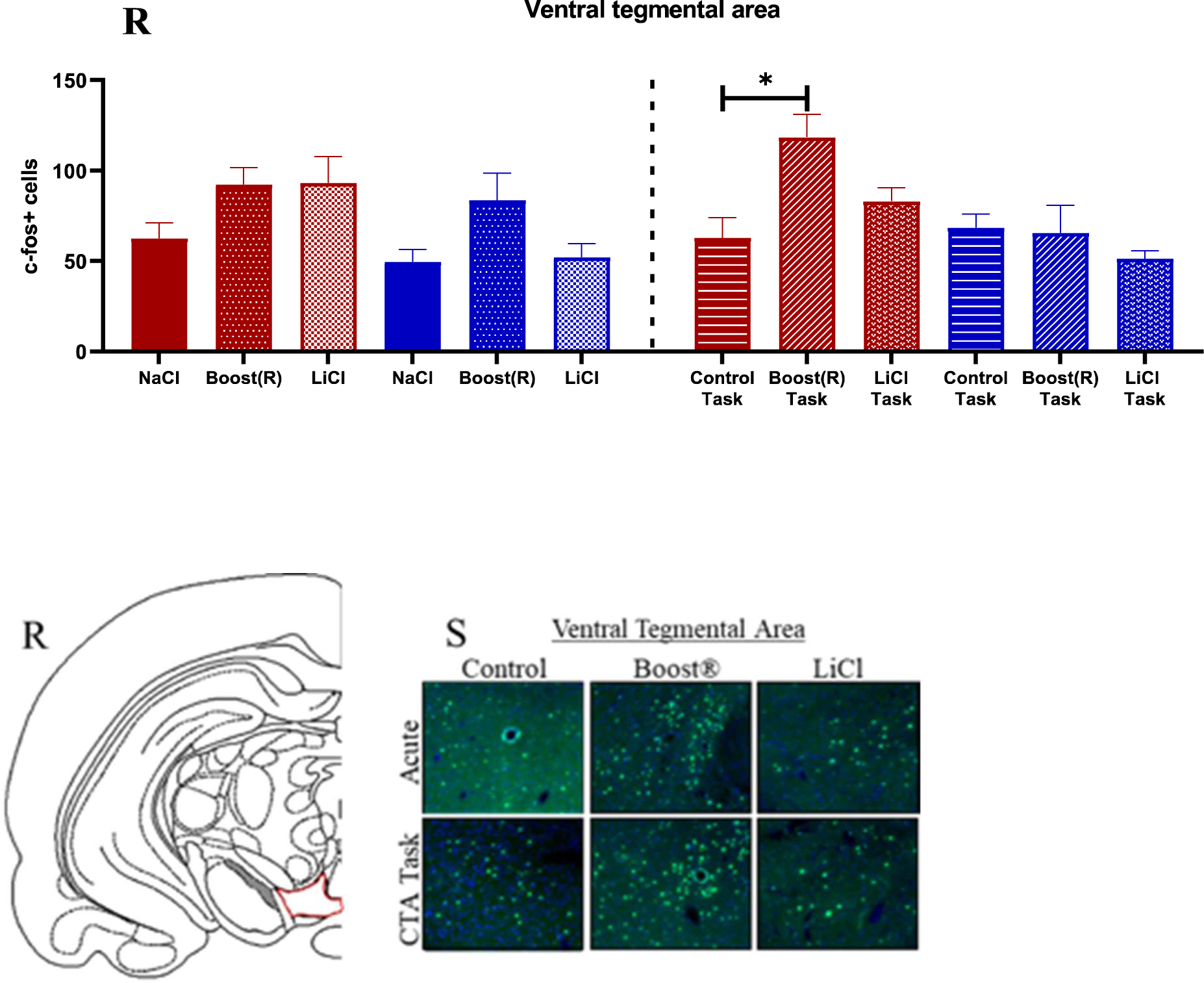
(A) c-fos response in the CeA across conditions. N=4-7. (B) c-fos response in the BLA across conditions. N=3-4 (C) Atlas image of CeA and BLA. (D-E), (F) c-Fos response in the gIC5/6 across conditions. N=4-9 (G) c-Fos response in the gIC4 across conditions. N=3-4 (H) Atlas image of gIC5/6 and gIC4. (J-K) Representative images. (L) c-Fos expression in the PVN across conditions. N=3-8. (M) c-Fos expression in the SON across conditions. N=3-4 (N) Atlas image of SON and PVN. (O-P) Representative images. (N) c-Fos response across conditions in the VTA. N=3-4 (Q) Atlas image of the VTA. (R) Representative images (S)

The granular insula (gIC) receives both taste and visceral inputs ^35–42^, and is divided into several layers. We considered layer IV (gIC4) and layers V/VI (gIC5/6). C-Fos in gIC5/6 increased with both Boost® and LiCl (Fig. 4f, atlas location shown in Fig. 4h representative images shown in Fig. 4j), showing a significant effect of treatment [gIC5/6; F (2,29) =14.52, p< 0.001]. c-Fos in gIC4 likewise increased in both Boost® and LiCl groups compared to NaCl control (Fig. 4g, atlas location shown in Fig. 4h representative images shown in Fig. 4k). The gIC4 showed an effect of treatment [F (2,17) =15.62, p> 0.001].

The supraoptic nucleus (SON) responds to acute LiCl ^43, 44^. It is the primary site of oxytocin production, which is known to have emetic and anorectic properties ^45–48^. The c-Fos response in the SON increased markedly in response to LiCl [F (2,16) =54.36, p< 0.001 effect of treatment], with no response to Boost® (Fig. 4l, atlas location shown in Fig. 4n, representative images shown in Fig. 4o). c-Fos in the paraventricular nucleus (PVN), which contains receptors for oxytocin and is the primary location for release of stress-related peptides such as corticotropin-releasing hormone and vasopressin ^49–52^, likewise increased specifically in response to the aversive stimulus [F(2,27)=8.44, p< 0.001, Fig. 4m, atlas image shown in Fig. 4n, representative images shown in Fig. 4p].

The ventral tegmental area (VTA) is responsive to rewarding stimuli and implicated in addiction ^53–55^. C-Fos in the VTA showed an effect of treatment [F (2,22) =4.19, p=0.03], with acute Boost® increased compared to acute LiCl and NaCl conditions (Fig. 4q, atlas image shown in Fig. 4r, representative images shown in Fig. 4s).

Interoceptive signals and contextual stimuli converge in the ventromedial frontal cortex (vmPFC) and agranular insula (aIC) ^39, 56–62^. However, no significant effects were found in the vmPFC or aIC in any condition, and so these will not be discussed further.

The nucleus accumbens responds to rewarding stimuli, typically in behavioral paradigms that utilize deprivation. The accumbens shell is more responsive to rewarding stimuli, while the accumbens core responds to conditioning of rewarding stimuli ^63–69^. No significant effects were found in either the accumbens core (NAcC) or shell (NAcS) during the acute stimuli.

### c-Fos immunoreactivity during CTA expression

3-way repeated measures ANOVA (sex x treatment x area) of the CTA task conditions revealed a main effect of sex [F (1,31) =7.33, p< 0.011] and area [F (10,231) =145.72, p< 0.001] and an interaction of treatment x area [F (20,231) =6.87, p=<0.001]. Females were more responsive than males overall. We then ran second order ANOVAs of treatment x sex in individual areas, and treatment x area in each sex.

c-Fos in the CeA increased in response to the Boost® task. It showed an effect of treatment [F (2,29) =24.95, p< 0.001] and just missed an effect of sex [F (1,29) =3.96, p< 0.056]. The Boost® task differed from the control task, both globally and in each sex individually (Fig. 4a, atlas location shown in Fig. 4c, representative images shown in Fig. 4d). c-Fos in the BLA showed an effect of sex [F (1,18) =17.34, p < 0.001, Fig. 4b, atlas location shown in Fig. 4c, representative images shown in Fig. 4e]. This was driven largely by high baseline c-Fos immunoreactivity in the control task for females. Lower-level ANOVA by sex showed a significant effect of condition in males only, with both Boost® task and LiCl task differing from control.

The gIC5/6 showed an effect of treatment [F (2,31) =5.29, p< 0.011], with the Boost® task differing from the control task. This difference is driven by the female rats, which showed increases in c-Fos response in both the Boost® task and LiCl task compared to control (Fig. 4f, atlas location shown in Fig. 4h, representative images shown in Fig. 4j). c-Fos in the gIC4 showed an effect of treatment [F (2,17) =4.39, p< 0.014], and no effect of sex (Fig. 4g, atlas location shown in Fig. 4c, representative images shown in Fig. 4k). The Boost® task showed increased c-Fos expression compared to the LiCl task globally. The females showed a statistically significant increase in c-Fos expression in the Boost® task compared to LiCl task conditions.

Females had higher c-Fos expression in the NAcC than males. It showed an effect of sex [F (1,31) =12.31, p< 0.001] but no effect of treatment (data not shown). Females had higher c-Fos expression than males, both globally and in the Boost® task specifically. c-Fos in the NAcS. showed an effect of sex [n=4, F (1,18) =14.52, p< 0.001], with no effect of treatment (data not shown).

Neither the PVN nor the SON responded during the task conditions. In the ventral tegmental area in the task conditions, there was an effect of treatment [F (2,16) =3.86, p=0.043], an effect of sex [F (1,16) =9.18, p=0.01], and an interaction of sex x treatment [F (2,16) =3.90, p=0.04]. By Fisher’s post hoc, females were increased compared to males overall, and Boost® task in females was increased compared to their control, as well as compared to males in the Boost® task (Fig. 4q).

### Network analysis of c-Fos responses to control, Boost®, and LiCl tasks

A major goal of the present study was to compare activated neural networks involved in the typical LiCl conditioned taste aversion task (LiCl task) with those activated when animals were anticipating Boost® presentation without a previous aversive stimulus (Boost® task) and those activated when animals experienced only the test cage (Control task). We first conducted network analysis of these three conditions.

We then subtracted correlations contained in the Control conditions from the Boost® task and LiCl task networks (Fig. 5a and Fig. 5b respectively). The two networks are quite distinct.

**Figure 5.**
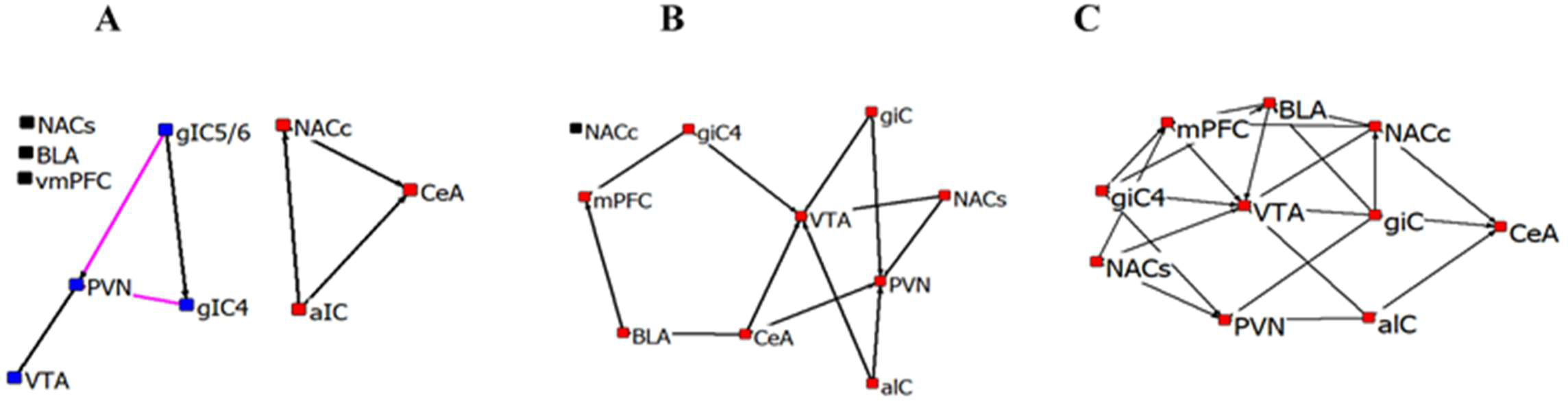
Network of significant c-Fos correlations for control task (A), Boost® Task-Control (B) and LiCl (CTA) Task-Control Task (C). Each brain region represented as a node in the network. Colors indicate correlations with no other brain areas (black), members of the main network (red) and members of a second network (blue, Control task). Line color indicates positive (black) or negative (magenta) correlations.

However in both, the VTA became central to each network, with all brain areas sample involved in the LiCl task-control task network. Responses of dopamine D1- and D2-receptor expressing cells in the amygdala to rewarding and aversive stimuli The role of the amygdala in the response to acute aversive stimuli such as LiCl is well-established ^43, 44, 65, 70–73^. A study by Kim, et. al., examined the expression patterns and responses to both fear and reward in multiple neuronal cell types in the CeA and BLA ^30^, laying an important groundwork for further interrogation into the function of specific neuronal cell types. The BLA is responsible for assessing the salience of a signal, whether it be rewarding or aversive^30–33, 74–76^. BLA outputs to the CeA promote defensive and appetitive behaviors ^30^.

Findings above indicating the centrality of the VTA in the task networks stimulated interest in potential amygdala dopamine involvements in this task. Dopamine receptors are present throughout the amygdala, but there is little known about their reactivity to reinforcing or aversive stimuli. We used RNAscope to visualize D1 and D2 dopamine receptors (Drd1 and Drd2) throughout the amygdala. We found that Drd1+ and Drd2+ cells occupy largely distinct anatomical locations within the amygdala. Consistent with previous studies, we found that the CeC and CeL were composed primarily of Drd2+ cells, while the CeM contained primarily Drd1+ cells. A smaller population of Drd1+ cells are present in the CeL (Fig 6a). The BLA, in contrast to the CeA, represents a nearly homogenous population of D1-receptor expressing cells, with few D2-receptor expressing cells (Fig. 6a).

**Figure 6.**
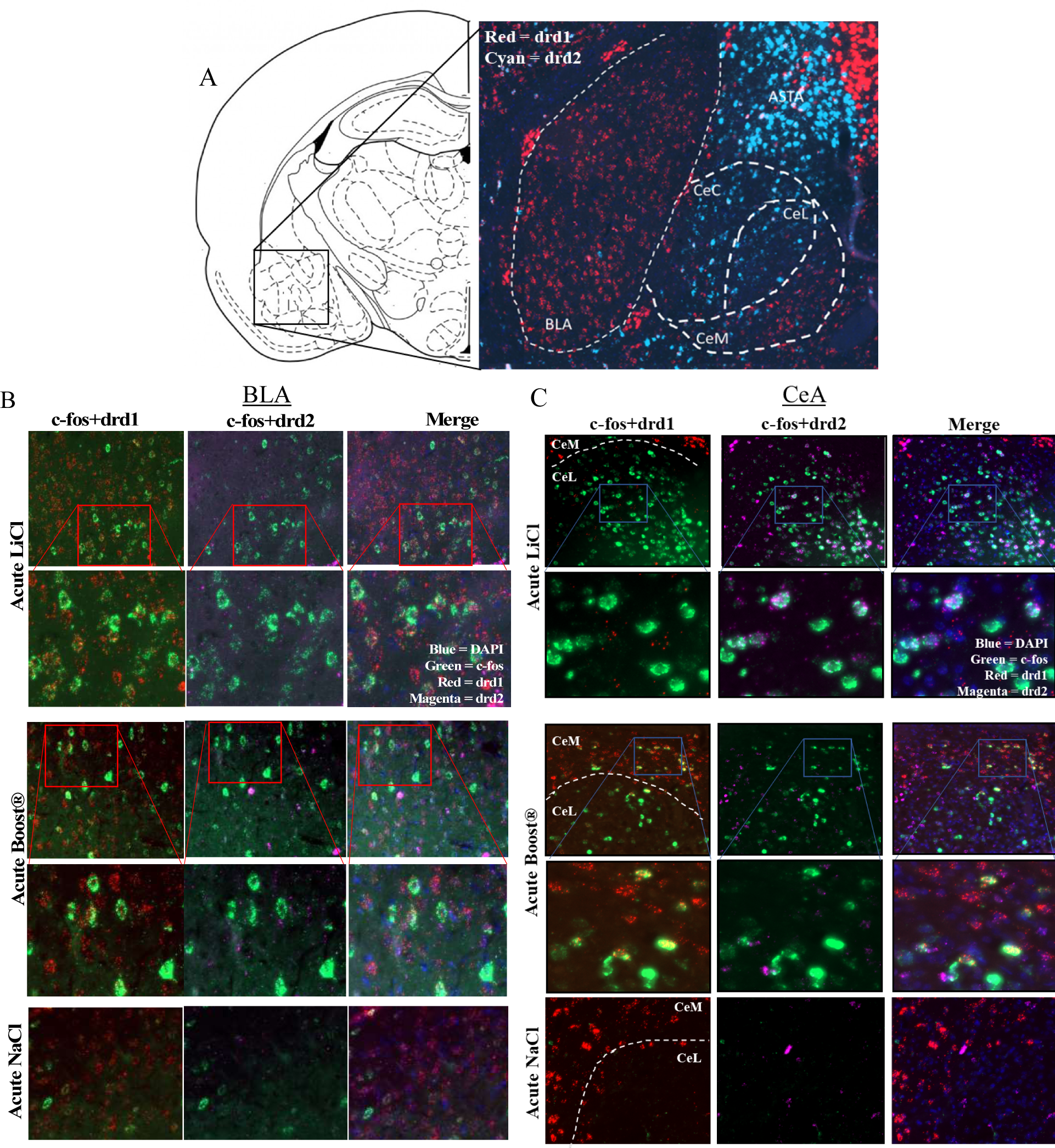
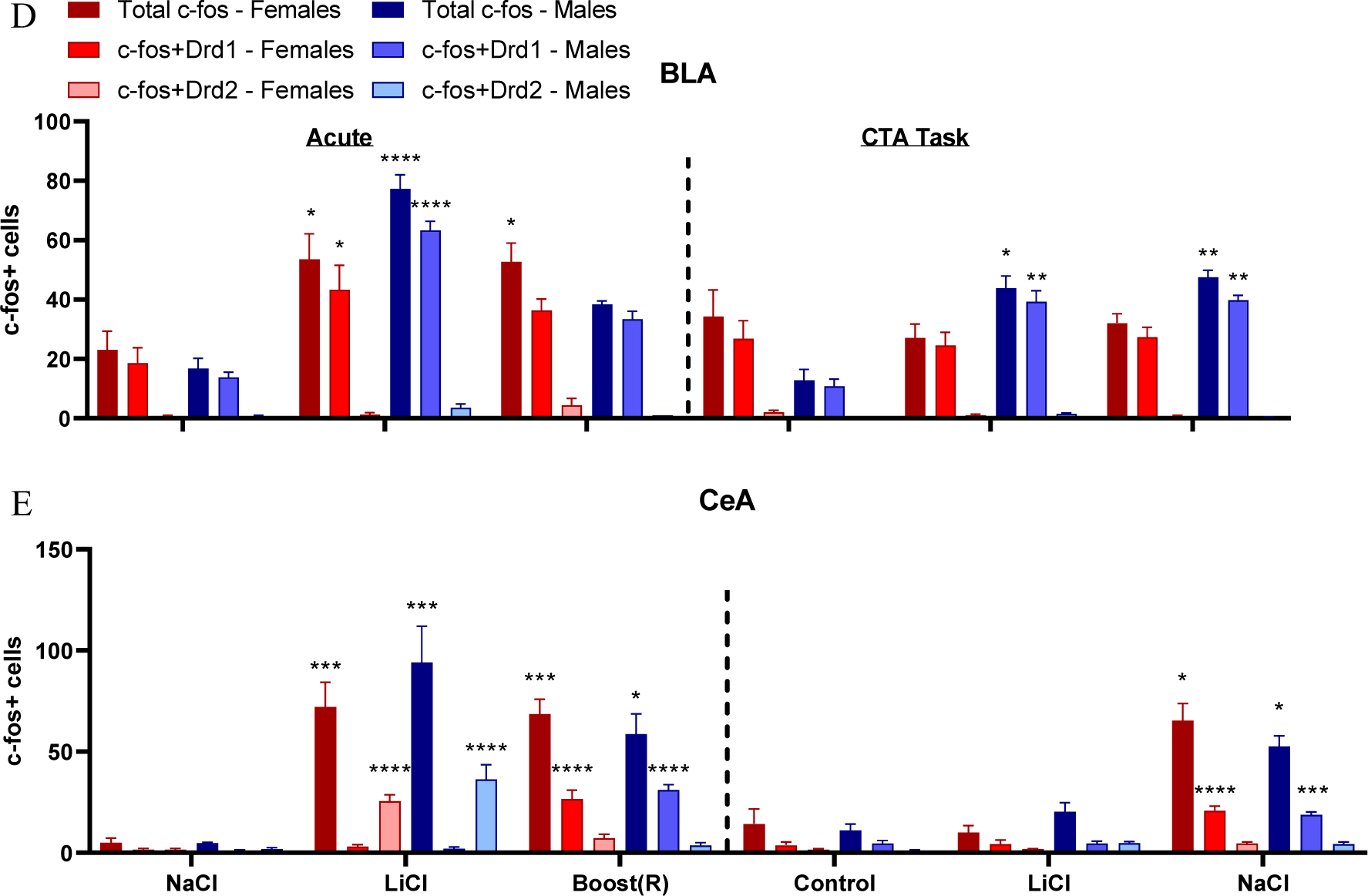
(A) Distribution of Drd1+ and Drd2+ cells across the amygdala. BLA = basolateral amygdala. CeA = central nucleus of the amygdala. ASTA = amygdalostriatal transition area. (B) Representative images of RNAScope in acute LiCl and acute Boost® ® conditions in the CeA. (C) Representative images of RNAScope in acute LiCl and Boost® ® conditions in the BLA. (D) RNAScope analysis of co-expression of c-Fos and Drd1+/Drd2+ cells in the basolateral amygdala across conditions. N=3-4/condition (E) RNAScope analysis of co-expression of c-Fos and Drd1+/Drd2+ cells in the central nucleus of the amygdala across conditions. N=3-4/condition.

Using the novel mRNA in situ hybridization method RNAScope, the co-expression of c-Fos and D1 or D2 receptors was characterized in each of the acute conditions, as well as during CTA expression, in the BLA and CeA. A representative image of the amygdala is shown in Fig. 6a.

### Responses of D1-receptor expressing cells in the BLA to acute and conditioned rewarding and aversive stimuli

Representative images for the BLA are shown in Fig. 6b and results in Fig. 6d. The BLA expressed c-Fos in response to both rewarding and aversive stimuli, and during CTA expression in male rats only. The Drd2 cells, which represented a small fraction of total cells, did not respond in any stimulus-specific pattern.

We performed 2-way ANOVA (sex x treatment) for total c-Fos mRNA. In the acute conditions, total c-Fos mRNA expression in the BLA showed a main effect of treatment [F (2,17) =32.00, p< 0.001] and an effect of treatment by sex [F (2,17) =5.73, p< 0.013] for the acute stimuli. Post hoc analysis showed Boost® and LiCl conditions differed from NaCl control, globally and for both sexes individually. We next considered the co-expression of c-Fos+/Drd1+ cells. 2-way ANOVA (sex x treatment) revealed an effect of treatment [F (2,22) =4.03, p< 0.005] and an interaction of treatment x sex [F (2,22) =4.03, p< 0.037]. Post hoc analyses revealed a difference between the acute Boost® and LiCl conditions compared to NaCl control, both globally and by each sex individually.

For the task condition, 2-way ANOVA revealed a main effect of treatment [F (2,23) =5.66, p< 0.001] and an interaction of sex x treatment [F (2,23) =9.46, p< 0.0016]. There was no main effect of sex. Post hoc analysis revealed both the Boost® and LiCl tasks differed from the control task. By sex, males in the Boost® and LiCl tasks differed from their respective controls. Females did not differ from controls, which had high levels of baseline c-Fos.

The same pattern emerged when considering c-Fos+/Drd1+ co-expressing cells. There was an effect of treatment [F (2,23) =8.59, p< 0.0024] and an interaction of sex x treatment [F (2,23) =9.64, p 0.0014]. There was no main effect of sex. As with total c-Fos, the Boost® and LiCl task conditions showed greater c-Fos mRNA expression than the control. As with total c-Fos only males differed in the Boost® and LiCl task conditions from their respective controls.

### The CeA is responsive to rewarding and aversive stimuli in a cell-specific manner

Representative images for the CeA are shown in Fig. 6c and results in Fig. 6e. Unlike the BLA, the CeA responded to rewarding and aversive stimuli in a cell-specific manner. In the acute condition, ANOVA revealed an effect of treatment [F (2,17) =30.32 p < 0.001] for total c-Fos+ cells. There was no main effect of sex or treatment x sex interaction. Post hoc analysis revealed the NaCl differed from both the Boost® and LiCl conditions. When considering c-Fos+/Drd1+ co-expressing cells, there was an effect of treatment [F(2,17) =100.22, p=< 0.001]. There was no main effect of sex or interaction of sex x treatment. The acute Boost® condition differed from the acute NaCl condition, with robust expression of double positive cells (c-Fos and Drd1). There was no difference between the acute LiCl and acute NaCl condition.

The opposite association of c-Fos and condition was found for Drd2 expressing cells. By ANOVA there was an effect of treatment [F (2,17) =38.63, p < 0.001]. There was no effect of sex or interaction of treatment x sex. The acute LiCl condition showed significant expression of c-Fos in Drd2+ cells, but not the acute Boost® condition.

In the task condition, there was an effect of treatment [F (2,18) =40.24, p=<0.001], with no effect of sex or treatment x sex interaction. There was significant mRNA expression of c-Fos in the Boost® task, but not the LiCl task condition. When considering c-Fos+/Drd1+ cells, there was an effect of treatment [F (2,18) =53.15, p < 0.001]. As with the acute condition, there was significant overlap between c-Fos+ and Drd1+ cells.

Altogether, these data showed that Drd1+ cells are differentially responsive to the rewarding stimulus in the CeA, while Drd2+ cells are differentially responsive to the aversive stimulus in the CeA. Following treatment with acute LiCl, c-Fos response showed both anatomical and neuronal cell type specificity, with activation occurring primarily in the CeL and CeC, and co-expressing with Drd2-expressing cells. Conversely, in c-Fos and c-Fos/drd1 co-expression were observed in both acute and conditioned Boost®.

Boost® produced a c-Fos response in the CeM and CeL and co-expressed with Drd1-expressing cells. Increases in c-Fos and c-Fos/drd1 co-expression were observed in both acute and conditioned Boost®.

## Discussion

Conditioned taste aversion is most often characterized as a task which reflects avoidance/aversion. By assessing behavioral and neural responses to both the reinforcing and aversive stimulus as well as the reinforcing and aversive task (Boost® task and LiCl task), the present findings reveal how animals balance the competing signals of safety or danger to learn and perform this task. Interrogation of the neural network revealed that stimuli activate circuits in a stimulus- and cell type-specific manner, suggesting that opposing stimuli activate parallel, but distinct, neural pathways.

### Behavioral responses to rewarding and aversive stimuli

CTA is an essential behavior to prevent further consumption of foods that may be toxic and should be avoided. The devaluation of the hedonic stimulus when it is paired with an aversive visceral stimulus is critical to the CTA paradigm. For example, delivering only an aversive stimulus like footshock can reduce CS consumption without changing palatability, while aversive visceral experience reduces both consumption and palatability, as measured by CS consumption in the home cage ^77–79^. Here, we use USVs to show that the behavior is more complex than simply showing aversion. Animals are both attaching a hedonic value to the rewarding stimulus and signaling the danger of the devalued stimulus.

This is the first study to comprehensively evaluate nausea behaviors in both male and female rats in CTA. We found that male and female rats showed similar nausea behaviors after LiCl administration. Of the three acute nausea behaviors we observed, only lying on belly correlated with CTA expression, suggesting this behavior may be an expression of more severe nausea than pica. Further studies in CTA and other nausea-relevant paradigms may prove useful in characterizing how this behavior correlates to nausea.

USVs were remarkably sex specific. They proved to be a useful tool to assess both positive (55 kHz) and negative (22 kHz) response to stimuli during CTA expression. Males expressed 55 kHz calls in anticipation of Boost® and inhibited 55 kHz calls when the Boost® was devalued with LiCl. The expression of USVs offers insight into the valuation animals are assigning to the Boost® and to the LiCl. Increases in 55 kHz in the context previously paired with Boost® suggests animals are assigning a positive valuation to the hedonic stimulus. The converse inhibition of 55 kHz USVs reinforces the notion that USVs can be correlated with an expected outcome. Vocalization patterns in females reflected high-estradiol (more vocal) and low-estradiol (less vocal) states but females were more likely to engage in 22 kHz calls than males. This result could suggest that females are more likely to attach a negative valuation to LiCl-paired Boost® or that they are more likely, in general, to elicit warning calls to alert the colony of nearby dangers. The latter hypothesis is supported by studies showing females are more likely than males to increase 22 kHz calls in the presence of a predator ^80^, especially considering their comparable sensitivity to CTA. USVs provide additional insight not only into the animal’s valuation of the stimulus, but to how they communicate this valuation to others in the colony.

### The c-Fos response to rewarding and aversive stimuli

CTA is a unique form of learning in that the stimuli do not need the same temporal proximity as other operant learning and behavioral conditioning paradigms ^81, 82^. The nature of digestion is such that visceral malaise may occur minutes to hours after the ingestion of a toxic substance. Hedonic and aversive inputs follow parallel pathways: caudal regions, such as the nucleus of the solitary tract and parabrachial nucleus, increase c-Fos in response to both LiCl and sucrose ^83, 84^. These circuits then converge in brain regions like the amygdala to assess valence.

We hypothesize intersection of the neural circuits conveying information about reinforcing and aversive stimuli may be found within the circuit itself, reducing the necessity for events to overlap in time for learning to occur.

This schema was borne out in our data. The neural circuit underlying the response to the acute stimuli overlapped in key brain regions, including the BLA, the CeA, the gIC5/6, and the gIC4. In contrast to brain areas that respond to both acute stimuli, the PVN and SON uniquely responded to the aversive stimuli, and the VTA responded uniquely to the reinforcing stimuli. Sampling of more caudal areas may prove useful in delineating the cell type-specific circuit in future studies. Regions such as the NTS ^85–92^ and PBN ^71, 72, 84, 86, 92, 93^ which were omitted in the present study to focus on rostral brain areas that are involved in higher-order processing of both aversive (emetic) and rewarding stimuli, and future use of cell-type specific probes will allow more complete dissection of this circuit .

The gustatory cortex (GC, which processes taste ^36, 37, 41, 94^) and visceral insular cortex (VIC, which processes visceral stimuli ^41, 42, 95, 96^) are topographically distributed throughout the insular cortex, including layers IV, V, and VI ^41, 42, 95, 97, 98^. Our data demonstrate that the GC and VIC show significant overlap in c-Fos expression ^41, 99^. The GC reacts to novel taste and lesions are known to cause a decrease in neophobia ^37, 94, 100, 101^. Additionally, lesions to the GC can attenuate CTA on first trial, a deficit that disappears with repeated CS-US pairings ^38^. These studies raise the question whether the loss of neophobia acts as a latent inhibition, reducing CTA by making the CS appear familiar ^102^. An alternative explanation from our studies is the possibility that lesions of the GC likewise destroy cells that are part of the VIC, and thereby reduce the nausea effects of LiCl. Lesion studies that also assess nausea behavior would help to clarify this question. Additionally, our findings of cell-specific c-Fos responses in the CeA suggested that future studies which interrogate cell-specific responses in areas like the highly heterogenous insular cortex to rewarding and aversive stimuli in a cell-specific manner will provide new insight into integration of affective valence to produce motivated behavior.

CTA is often characterized as analogous to fear learning. However, CTA involves the convergence of two stimuli of opposing valence, resulting in complex decision-making rather than a fear-induced defensive response. The RNAScope data showing that the response to rewarding and aversive stimuli show cell-type and anatomical specificity in the CeA suggests that other regions may show similar stimulus-specificity when specific cell populations are evaluated. How these two neural networks converge to produce the memory of the experience, and the subsequent decision-making process that guides the animal’s behavior, is an important area of future study.

We were surprised to see there was no activation of the nucleus accumbens shell after acute Boost®, or the core during expression of the task. One explanation is that most studies showing such activation increased the reward of the hedonic substance by water and/or food deprivation during CTA acquisition, while we did our studies in animals that were not deprived. Studies show deprived animals express higher levels of c-Fos in these key brain areas compared to satiated animals ^66, 68^.

We found no sex differences in the neural activation in response to the acute stimuli in any of 11 brain regions we assessed. This was a surprising result, as females drink more Boost® at first exposure and are relatively less sensitive to developing CTA. It is possible that our limited analysis did not sample a critical area that accounts for these differences. A more comprehensive study of the neural circuit that includes other brain regions associated with CTA, such as the bed nucleus of the stria terminalis or parabrachial nucleus may be of value.

### The c-Fos response during CTA expression

We used a novel experimental paradigm for studying the neural circuits engaged during the acute rewarding and aversive stimuli, as well as during the reinforcing task (expectation of Boost®) and CTA task (expectation of Boost® after pairing with LiCl). Neural activation during the Boost® task was similar to that of the acute Boost®, while activation during the LiCl task differed considerably from its acute stimulus. The rewarding effects of Boost® that were not devalued with LiCl are reinforced with repeated offerings of the hedonic stimulus. We therefore saw similar neural activation during expression of the Boost® task. There is no increase in c-Fos with the second pairing, likely because the taste is no longer novel ^103, 104^. When this same stimulus was devalued with LiCl, drinking behavior was reduced, and the circuit was inhibited.

During CTA expression, females broadly expressed higher levels of c-Fos than males. By area, they were significantly more responsive in the gIC5/6 and gIC4 during the Boost® task.

We also found that the VTA increased with the reinforcer (Boost® task) in only female rats. The VTA is known for its role in reinforcement and addiction, and sends outputs to the nucleus accumbens, amygdala, and frontal cortex ^54, 105–107^. Our data support the hypothesis that female rats find the sweet taste more reinforcing and are therefore less sensitive to CTA at lower doses of LiCl. Cell-type analysis would benefit our understanding of how females differentially process sweet taste, a critical area of study in the field of eating disorders ^108–116^. This result is particularly interesting considering their relative lack of USV expression in response to the rewarding stimulus. While males are more unequivocal about the “goodness” of Boost®, as evidenced by their consistent expression of 55 kHz USVs, females in low-estradiol states do not express 55 kHz USVs, despite drinking comparably.

### Network analysis of c-Fos correlations

Network analysis of c-Fos regional correlations during Control, Boost® and LiCl tasks provided insights into the c-Fos responses that were not revealed by simple analysis of c-Fos responses by condition. Each of the three task conditions elicited unique networks of brain area interactions. The control task was the simplest, with two independent small networks and 3 areas with no interactions (vmPFC, NACs and BLA). In the Boost® task and the LiCl task, the aIC was a central node in the more complex interactions, a finding consistent with its known role in ingestive tasks and suppression of ingestion associated with sickness^117^. The BLA exhibited correlated responses with aIC and vmPFC in both tasks, also a relationship predicted by BLA’s known role in learned responses (including to taste and ingested stimuli), both appetitive and aversive^75, 118, 119^.

The most impressive demonstration from this analysis is the centrality of the VTA in both Boost® -Control and LiCl-Task networks, and its correlations with PFC, NACs, BLA and CeA with the Boost® task and both NACs and NACc in the LiCl-Control task. A significant body of work has identified specific subpopulations of dopamine neurons in the VTA that respond to positive and negative stimuli through interaction with nucleus accumbens and prefrontal cortex, as well as its innervation of the CeA and BLA^117, 120–124^. All these associations are present in this single model.

Network analysis also revealed previously unexpected relationships. The presence of the PVN and vmPFC in all three task networks in both Boost® and LiCl was not predicted from their c-Fos levels in individual tasks, given their lack of significant c-Fos response to task. For PVN, its conflicting negative relationship at least with gIC5/6 and gIC4 suggests that heterogenous populations in the brain area were activated by varying aspects of these conditions. This speculation is strengthened by the more central involvement of the PVN in the Boost® -

Control and LiCl-Control task networks once these Control influences were subtracted. vmPFC c-Fos was the highest in almost all conditions, but the specificity of its role in each condition might not have been revealed without network analysis.

The multiple interactions revealed from the modest number of brain areas sampled (11) demonstrates the potential value of network analysis, especially the subtraction approach of removing correlations associated with control conditions. For example, it suggests that mechanistic interrogation of dopaminergic influences in CeA and BLA during CTA with approaches like optogenetics and/or DREADDS could be informative. More complex networks can be identified by sampling many brain areas, as has been shown for fear memory^125^ and in response to both approach and avoidance associated with social interactions^126^.

### Dopamine receptors in the amygdala respond to rewarding and aversive stimuli

Our studies showed that Drd1+ neurons were responsive to the rewarding stimulus, while Drd2+ cells were responsive to the aversive stimulus. This is the first time these cell types have been studied in the context of CTA. Kim, et. al. ^30^ showed that silencing Drd1+ neurons in the CeA inhibited feeding behavior in mice. However, other studies demonstrate that direct blockade of Drd1 in the CeA did not affect feeding behavior, suggesting these neurons are being activated by something other than dopamine. Drd1+ neurons highly co-express with 3 other neuropeptides: tachykinin 2, neurotensin, and somatostatin. Although these neuronal cell types do not affect behavior when manipulated on their own, Kim shows that these cell types may work collectively to coordinate feeding behavior.

Only about 30% of c-Fos+ neurons in the CeA co-expressed with Drd1+ cells in response to the appetitive stimuli, suggesting additional cell types may also be important for the processing of the reward. A similar percentage of Drd2+ receptors were responsive to LiCl. A more thorough examination of the CeA in response to a food reward and aversive stimulus would be useful in delineating the cell types that process this stimulus.

Contrary to our findings in the CeA, Drd1+ neurons in the BLA responded to both rewarding and aversive stimuli, without input from Drd2+ neurons. These findings suggest that Drd1+ neurons may receive innervation from more caudal areas that respond specifically to these stimuli ^127^. Additionally, the BLA is known to respond to rewarding or aversive stimuli in a location-specific manner, with the rostral BLA responding to aversive stimuli and the caudal BLA responding to rewarding stimuli. The dual responses observed in the present study may reflect the location at which we sampled the BLA, which was in the central zone where these two stimuli converge ^128^.

In summary, these findings support our hypothesis that CTA is best characterized as a task in which animals must balance rewarding and aversive stimuli, and therefore is not simply a model of fear learning ^129–133^. As such, it has significant relevance as a task which reveals how affective valence is integrated into decisions about behavior. The USV analysis showing inhibition of 55 kHz calls and increased 22 kHz calls in the LiCl-paired context suggests that additional neural centers may be involved in assigning valence during the CTA task that are not captured by LiCl behaviors or our c-Fos analysis. Our c-Fos analysis of rostral brain regions revealed a significant role of the CeA and granular insula, which represents both the gustatory and viscerotopic cortices, in processing of the reinforcing and aversive stimuli and showed increased global c-Fos expression in females overall during the task. Further characterization of both the acute and conditioned circuits in males and females will help clarify how these stimuli are differentially processed between the sexes. Finally, our data found a novel role for specific cell types in the CeA and BLA that have not previously been studied. These results suggest that similar cell-specific analysis in cortical areas will continue to reveal details of the neural circuit(s) engaged while animals learn this type of task.

## Materials and Methods

### Animals

Male and female Sprague Dawley rats [post-natal day (PN) 60, Charles River Laboratories, Raleigh, NC] were received 1 week before behavior tests. Rats were housed by sex in ventilated plastic cages with ad libitum PM5001 Rodent Chow and water on a 7 am-7pm light/dark cycle. Females were not selected based on estrous cycle stage, but cycle was determined by lavage on the final day of behavioral testing. A single lavage was performed as repeated lavages can establish place preference for the location of lavage and suppress behavioral responses to other reinforcers ^134^.

### Conditioned Taste Aversion Protocol

Animals were conditioned to ip injections with NaCl for 3-4 consecutive days prior to experimental procedures. The conditioned taste aversion protocol (Fig. 1a) was conducted over 3 days. On day 0, animals were water deprived overnight and then conditioned to test cage. Water bottles were weighed and placed on cages with spouts pointing way. Rats were allowed to habituate to their new environment for 30 minutes, after which they were given access to water for 60 minutes in the test cage, to condition them to drinking from new bottles. Water bottles were weighed again, and consumption was measured.

On day 1, animals were returned to the test cage and allowed to habituate for 30 minutes. They were then offered Boost® and allowed to drink for 60 minutes. Boost® consumption was measured. If animals drank a minimum of 1 mL of Boost®, they were then injected with LiCl (38 mg/kg, 0.15M) or the equivalent volume of isotonic NaCl. This dose was selected to achieve a robust c-Fos response for interrogation of the neural network. They were observed for 60 minutes for LiCl-related behaviors. They were then returned to their home cages and allowed to drink and eat ad libitum.

On day 2, non-water deprived animals were returned to the same test cage and allowed to habituate for 30 minutes. They were then given a 2-bottle test, offering either Boost® or water for 60 minutes. Boost® and water consumption were measured, and the ratio of day 2/day 1 (D2/D1) consumption was calculated. After 60 minutes of re-exposure to the Boost®, animals were perfused with formalin and brains collected for c-Fos analysis. Brains were post-fixed in formalin overnight at 4°C, then cryoprotected in 30% sucrose solution (30% sucrose, 70% PBS) for 3-4 days. Brains were flash-frozen in brain molds in ethanol/dry ice bath, in 2:1 30% sucrose/tissue freezing media solution. They were stored at -80°C until ready to cut for immunohistochemistry or RNAScope.

Animals whose brains were analyzed for the acute stimulus were taken through day 0 as described. On day 1, they were allowed to habituate for 30 minutes in the test cage before receiving LiCl (38 mg/kg) or the equivalent of isotonic NaCl or were allowed to drink Boost® for one hour, after which they were perfused, and brains collected for c-Fos analysis (see Table 2).

**Table 2.** Experimental Conditions

| *Acute Stimulus* | *Conditioned Stimulus* |
| --- | --- |
| Boost® | Boost® paired with NaCl |
| LiCl (38 mg/kg) | Boost® paired with LiCl |
| NaCl (control) | Context only control |

### Behavior Scoring

Three behaviors were time-sampled as a behavioral estimation of nausea [pica, ptosis, lying on belly (LOB)]. As rats do not vomit, they will often attempt to dilute an ingested toxin through pica, the consumption of non-food substances. LiCl behaviors were assessed using a modified version of published measures: Pica and LOB are well known LiCl behaviors ^135–137^, and ptosis was included following the observation that it occurred frequently in LiCl-treated animals compared to NaCl-treated animals. Animals are observed for two consecutive 15-second blocks (30 seconds total) every 5 minutes for 1 hour. Animals were given a score of 0-3 as described in table 3.

**Table 3.** Scoring of LiCl Behaviors

| **Behavior** | **Score = 0** | **Score = 1** | **Score = 2** | **Score = 3** |
| --- | --- | --- | --- | --- |
| Pica | No pica | Some pica while still moving around | Pica in a crouched position, but still moved or looked around | Remained crouched, engaged in pica for most of 15-second interval |
| LOB | No LOB | Some LOB, still in crouched position | Some LOB, more flattened posture | Entirely flattened posture |
| Ptosis | No ptosis | Ptosis for <5 seconds | Ptosis for most interval (>5 seconds, <10 seconds) | Ptosis for entire interval |

### Scoring of Ultrasonic Vocalizations

In a within-groups design, rats were split by sex, treatment [NaCl + H2O (Control), NaCl + Boost®, and LiCl + Boost®] on Day 2 of CTA experiment (task day). USVs were recorded with an Echo Meter Touch 2, from Wildlife Acoustics, an accessory that attaches to a lightning port on the Apple iPhone. The Echo Meter Touch Bat Detector iOS app (v 2.7.20) was used to record the vocalizations. Recordings were uploaded from the smartphone to a secure online server. Manual scoring of the USVs during the 10-minute recording period was done using the visualization software from Wildlife Acoustics, Kaleidoscope Pro Analysis. Scoring was performed by a blinded observer in batches of 1-minute intervals. The scorer determined how often the rat engaged in >10 55 kHz vocalizations, or >1 22 kHz vocalizations in each interval. Scores were calculated as a percentage of the 10 1-minute blocks in which the vocalizations occurred.

### Immunohistochemistry

Brains were cut on a Leica CM3050S cryostat at 30 uM and stored in 1:1 TBS/glycerol solution at -20℃ until ready to stain. To stain, slices were first washed in 0.2% Triton-X in TBS solution 3 times. Slices were then blocked for 1 hour in solution of 0.3% Triton-X and 5% normal goat serum (NGS) in PBS. Slices were stained overnight in 0.3% Triton-X, 5% NGS, and 1:20,000 anti-c-Fos antibody (Abcam - ab190289) with gentle shaking at 4℃. Slices were then washed 3 times in solution of 0.3% Triton-X and 5% NGS in PBS (10 minutes first wash, 30 minutes second wash, 40 minutes third wash). Slices were then stained with secondary antibody (AlexaFluor 488, Invitrogen – A-11034) in 0.3% Triton-X, 5% NGS, and 1:200 secondary antibody for 2 hours. Slices were then washed in 1X PBS (10 minutes first wash, 30 minutes second wash, and 40 minutes third wash), with DAPI (R&D Systems 5748) added at a dilution of 1:10,000 for last 10 minutes of second wash. Slices were mounted on VWR Superfrost® Plus microscope slides with a drop of Vecta-Shield anti-fade medium (Vector Laboratories 101098- 042).

### Vaginal lavage and estrus cycle analysis

A vaginal lavage smear was performed on all female test subjects prior to transcardial perfusion. Vaginal cytology was analyzed to determine the phase of estrous cycle. The method of estrous cycle analysis as previously described ^134^. Animals were grouped into high estradiol states (proestrus/estrus) and low estradiol states (metestrus/diestrus).

### RNAScope

Brains were cut at 14 uM on a Leica CM3050S cryostat and placed on a Superfrost® Plus Micro Slide. 3 consecutive slices were placed on each slide. Slices were collected ∼90 uM apart. Slides were desiccated at -20°C for 20 minutes, then placed into -80°C with desiccants for long term storage. Slides were stained within 1 month of collection for optimal signal.

When ready to stain, slides were placed in 1X PBS for 5 minutes to remove OTC then placed in the HybEZ™ oven for 30 minutes at 60°C. They were then transferred to cold PFA for 15 minutes at 4°C, after which they were taken through the RNAScope procedure as outlined by ACDBio, using Protease IV. Probes were for c-Fos (Rn-Fos - 403591), Drd1 (Rn-Drd1a-C2, 317031-C2), and Drd2 (Rn-Drd2-C3 - 315641-C3). The Fos probe contained 20 oligo pairs and targeted region 84-1218 (Acc. No. NM_001256509.1) of the Fos transcript. The Drd1 probe contained 20 oligo pairs and targeted region 104-1053 (Acc. No. NM_012546.2) of the DRD1 transcript. The Drd2 probe contained 20 oligo pairs and targeted region 445-1531 (Acc. No. NM_012547.1) of the DRD2 transcript. Secondary antibodies were obtained from Akoya (OPAL 690, OPAL 520, and OPAL 570) and diluted to a concentration of 1:750. All experiments were run with positive and negative controls. The positive control targeted Polr2a (C1), PPIB (C2), and UBC (C3). The negative control targeted DapB (of Bacillus subtilis strain).

### Imaging and analysis

Slides were imaged with an Axio Imager upright microscope at 20x (16 z-stacks 2 uM apart) for IHC or 20x (8 z-stacks 1 uM apart) for RNAScope. Images were z-projected for max intensity using FIJI (ImageJ) software and c-Fos+ neurons were manually counted. 2-6 slices per region were imaged and counted per animal. Counts were averaged per animal.

### Drugs

LiCl was purchased from Sigma Aldrich (203637). LiCl was dissolved in distilled sterile water to a concentration of 0.15M. Sterile isotonic saline (NaCl, 0.9%; 0.15M) was used as a control injection.

### Network analysis

Network analysis of the Control task, Boost® task and LiCl task were conducted using slight modifications from a similar approach used to analyze networks activated by D1 and D2 agonists in developing rat brain^138^. Correlations among c-Fos levels in brain areas in each task condition were established by developing a matrix of correlations for each pair of brain areas for every subject. Males and females were combined to generate adequate statistical power to conduct the analysis. A symmetrical matrix of statistically significant (p < 0.05 or better) Pearson correlation coefficients (r) was created in NCSS. All other correlations were set at 0. This matrix was loaded into UCINET 6.370 (Analytic Technologies, Lexington KY)^139^. Networks for Control task, Boost® task and LiCl task were visualized with Netdraw. Each brain area appears as a node, and statistically significant correlations between specific pairs of brain areas are indicated as a line between brain areas. Figures are derived from UCINET graph theoretic layout with distance and node repulsion which provided the clearest illustration of all the correlations detected. Members of independent substructures identified by UCINET software (in Control Task only) are indicated by the color of the symbols. Brain areas with no c-Fos correlations with other brain areas are shown in the upper left. A secondary analysis of Boost® Task and LiCl task networks after subtraction of correlations associated with the Control task was conducted. All Pearson correlation coefficients were converted to Z scores using the Fischer transformation (1/2[ln (1 + r) – ln (1-r]) then Z scores for control conditions were subtracted from Boost® task and from LiCl task, Z-scores were back-translated to Pearson correlation coefficients, t values calculated using the formula r*√(n-2)/√(1-r^2^). Significant correlations in the subtraction networks were visualized as described above. This approach was selected in lieu of more typical heatmaps of Pearson correlation coefficients as it provides more information about relationships among multiple areas and is more rigorous because it includes only statistically significant correlations.

### Statistics

All results were analyzed by ANOVA with post hoc Fisher’s exact test corrected for multiple comparisons using the statistical package NCSS. Behavioral results were analyzed by 2- way ANOVA (sex x treatment). c-Fos responses were analyzed by 3-way repeated measures ANOVA (sex and condition as between measures and brain area as repeated measure). All conditions yielded area as a main effect, and second level ANOVAs were run for each brain area independently. Fisher’s LSD post hoc was used to determine groups that differed from control.

Tukey’s post hoc was used in RNAScope experiment. N for all behavior and c-Fos experiments was estimated by power analysis and variance observed in previous experiments with CTA behavior and c-Fos analysis of multiple brain regions ^85–87, 89, 140–143^. Individual differences shown in estrous cycle and USVs to illustrate individual variability in behavior.

